# Structural Connectome Analysis using a Graph-based Deep Model for Prediction of Non-Imaging Variables

**DOI:** 10.1101/2025.03.09.642165

**Authors:** Anees Kazi, Jocelyn Mora, Bruce Fischl, Adrian V. Dalca, Iman Aganj

## Abstract

We address the prediction of non-imaging variables based on structural brain connectivity derived from diffusion magnetic resonance images, using graph-based machine learning. We predict age and the mini-mental state examination (MMSE) score as examples of a demographic and a clinical variable. We propose a machine-learning model inspired by graph convolutional networks (GCNs), which takes a brain connectivity graph as input and processes the data separately through a parallel GCN mechanism with multiple branches. The novelty of our work lies in the model architecture, especially the Connectivity Attention Block, which learns an embedding representation of brain graphs while providing graph-level attention. We show experiments on publicly available datasets of PREVENT-AD and OASIS3. We validate our model by comparing it to existing methods and via ablations. This quantifies the degree to which the connectome varies depending on the task, which is important for improving our understanding of health and disease across the population. The proposed model generally demonstrates higher performance especially for age prediction compared to the existing machine-learning algorithms we tested, including classical methods and (graph and non-graph) deep learning.

## 2 Introduction

A structural connectome is a comprehensive map of the physical (anatomical) connections between different regions of the brain. It represents the white matter pathways that link various brain regions, forming a network that supports communication and information flow across the brain (Hofman, 2015; Sporns et al., 2005). Analyzing the structural connectome provides insight into neural circuitry, connectivity patterns, and their implications for cognition and behavior, contributing to our understanding of brain development, aging, and the impact of neurological conditions on brain wiring (Hagmann et al., 2008; F. Zhang et al., 2022). Connectomic analysis is a relatively new avenue for studying the brain, which, while not yet included in clinical practice, holds promise for neuroscientific discoveries and biomarker development. Given the complexity and the vast amount of data involved, deep learning (DL) techniques are likely to be effective in analyzing the connectome. Structural brain connectivity analysis is a data-driven approach that explores the overall structural connections in the brain. It involves studying patterns of anatomical connectivity via fiber tracking (tractography) throughout the brain (Zalesky et al., 2012).

Structural connectome graphs have been used to study a wide range of neurological and psychiatric disorders, including Alzheimer’s disease (AD) (Aganj et al., 2023; Amoroso et al., 2017; Frau-Pascual et al., 2021; He et al., 2008; J. Wang et al., 2018), schizophrenia (Karlsgodt et al., 2010; Shi et al., 2012; Y.-m. Wang et al., 2020), autism spectrum disorders (ASD) (Tolan & Isik, 2018), and Parkinson’s disease (Arrigo et al., 2019; Jellinger, 2022; X. Zhang et al., 2018; Y. Zhang et al., 2019), as well as to understand cognitive decline (X. Xu et al., 2021) and normal brain development and aging (Coelho et al., 2021; Damoiseaux, 2017; Dennis et al., 2013; Lewis et al., 2022; Neudorf et al., 2024). Numerous studies have identified specific structural and functional changes in the brains of patients with AD, such as altered functional activation, decreased volume or thickness of gray matter in certain brain areas, and abnormalities in the connections between different regions (Xie & He, 2012).

Structural brain connectivity analysis using artificial intelligence (AI) is an emerging field (Dubost, 2020; Sjöblom et al., 2020). The exploration of AI applications in deciphering dMRI data to study brain connectivity is an evolving domain that needs further investigation (Faiyaz et al., 2023). In this paper, we develop a technique to analyze dMRI-derived connectivity using DL for age and dementia prediction. Recently, Prescott et al. (2022) have shown that structural connectivity is a crucial factor in identifying early-onset AD risk, as individuals with a genetic predisposition show lower connectivity, especially in the frontoparietal control network, and this reduced connectivity is linked to the estimated time until dementia symptoms emerge. Toward this goal, one of the key tasks we address is predicting the Mini-Mental State Examination (MMSE) score of subjects directly from their structural connectomes using our proposed AI-based framework, thereby enabling quantitative estimation of cognitive decline associated with dementia.

Further, studying brain aging through the structural connectome is crucial as it provides insight into how brain connectivity and network organization change over time, which can be related to cognitive decline, neurodegenerative diseases, and overall brain health (Meunier et al., 2009). Accurate brain age prediction also serves as a potential biomarker for early detection of atypical aging trajectories, helping to identify individuals at risk of conditions such as AD and other forms of dementia. The task of predicting clinical and demographic variables from structural connectomic data, however, poses significant challenges due to the high dimensionality and non-Euclidean nature of brain connectivity. A recent work (Joo et al., 2023) shows that convolutional neural networks show suboptimal performance when used standalone, compared to using them in combination with Multi Layered Perceptrons (MLPs). Traditional DL approaches for age prediction have predominantly utilized convolutional neural networks (CNNs) applied to structural MRI data. A recent work (Peng et al., 2023) proposed a DL algorithm that leverages brain structural imaging data and enhances prediction accuracy by integrating biological sex information, employing a dataset of 3004 healthy subjects aged 18 and above using CNN algorithms on T1-weighted images. Similarly, lightweight CNN architectures such as Simple Fully Convolutional Networks (SFCN) have demonstrated accurate disease prediction with neuroimaging data, with CNNs showing the ability to accurately predict chronological age in healthy individuals from structural MRI brain scans (Cole et al., 2022; Peng et al., 2021). Machine learning (ML) algorithms trained to estimate age from brain structural MRI using DL-based approaches have revealed genetic associations, with new DL methods uncovering associated sequence variants (Cole et al., 2019). Recent advances include anatomical feature attention-enhanced 3D-CNNs for age prediction from structural MRI data, with CNN architectures capturing structural feature changes of brain aging based on MRI to predict age in healthy individuals (Feng et al., 2024; Zhao et al., 2024).

DL approaches for MMSE prediction have been less extensively explored compared to age prediction, with most studies focusing on joint prediction tasks combining disease diagnosis and cognitive assessment. Ryu et al. (2019) developed a 3D CNN architecture for both AD classification and joint MMSE score prediction using resting-state functional MRI scans of 331 participants, obtaining functional 3-dimensional independent component spatial maps as features for both classification and regression tasks (Ryu et al., 2019). The limited number of studies specifically targeting MMSE prediction using DL suggests this remains an underexplored area, with most cognitive assessment predictions relying on multimodal approaches or functional connectivity data rather than structural MRI alone.

Graph neural networks (GNNs), particularly the graph convolutional network (GCN) architecture (S. Zhang et al., 2019), provide a powerful and flexible framework for analyzing brain connectivity data. GCNs operate on graph representations of the brain, where nodes correspond to anatomical regions and edges encode structural connectivity (e.g., fiber density or tract strength), enabling the model to integrate information from each region along with the context provided by its connected neighbors. Thus, GCNs leverage the rich structural information in the connectome to make accurate predictions. They do so by performing iterative message-passing between neighboring nodes in the graph, using learnable functions to aggregate and transform information from neighboring nodes, and updating the features of each node based on the aggregated information. This allows GCNs to capture the complex relationships between brain regions and their connections, and make predictions based on this information. A GCN-based model is promising for analyzing the structural connectome and has shown great potential for improving our understanding of neurological and psychiatric disorders (Kazi, Shekarforoush, Arvind Krishna, et al., 2019; Kazi, Shekarforoush, Kortuem, et al., 2019), as well as normal brain development and aging (X. Li et al., 2020).

Recent GCNs have been employed for functional brain network analysis (X. Li et al., 2021), psychiatric disorder diagnosis (Zheng et al., 2024), optimization of GNN architectures for schizophrenia spectrum disorder prediction (S. Wang et al., 2024), classification of ASD versus hyper complex brain networks (Hu et al., 2021), and sex classification (Ktena et al., 2018). Many papers have shown work using structural brain networks for causal inference (Wein et al., 2021), brain age estimation (Moon et al., 2024), early diagnosis of AD (Y. Zhang, He, et al., 2023), and schizophrenia diagnosis (Sebenius et al., 2021), achieving high performance compared to existing ML, including DL, methods. GCNs have also been combined with recurrent neural networks to predict sex on temporal functional brain connectivity graphs (Kazi et al., 2022). A spectral GCN has been employed for region-of-interest identification in functional connectivity graphs and for sex classification (Arslan et al., 2018). In a multi-modal integration technique, Qu et al., 2025, the authors proposed a MaskGNN framework that integrates multimodal brain connectivity data for predictive analysis. Specifically, functional connectivity (FC) from fMRI and structural connectivity (SC) from DTI were combined at the node level and used as inputs to the MaskGNN model. In the latent space, the model further incorporated anatomical statistics (AS) from sMRI and computed structure–function coupling to capture complementary relationships between modalities. The aggregated features were processed through MaskGNN embedding, pooling, and readout layers, enabling effective multimodal feature learning.

Methodologically, most GCN methods use the Laplacian-filter based spectral implementation (Kipf & Welling, 2016). GCNs suffer from limited generalization to new graphs due to their dependency on the fixed graph Laplacian (Q. Li et al., 2018; Oono & Suzuki, 2019), and they struggle with over-smoothing in deep architectures where node representations become indistinguishable. Graph attention networks (GATs) (Veličkovíc et al., 2017) incorporate attention mechanisms for adaptive edge weighting, enabling nodes to selectively attend to their neighborhood features without requiring costly matrix operations or prior knowledge of graph structure. GATs leverage masked self-attentional layers to address shortcomings of prior graph convolution methods by enabling nodes to attend over their neighborhoods’ features and implicitly specify different weights to different nodes in a neighborhood without requiring costly matrix operations or depending on knowing the graph structure upfront. Despite these advantages, GATs are computationally inefficient for capturing long-range dependencies due to over-smoothing issues (Dasoulas et al., 2021) and exhibit high computational costs when applied to dense graphs (Wu et al., 2020).

Dynamic Graph CNNs (DGCNNs) (Phan et al., 2018) are employed for temporal graph analysis of multi-level graph structures and enable hierarchical learning through spatiotemporal integration of spatial and temporal data. While DGCNNs can adapt to changing graph topologies over time, they suffer from computational inefficiency and instability due to repeated graph construction at each layer, leading to high memory usage and potential noise amplification. Both GCN and GAT methods face scalability challenges with large graphs (S. Liu et al., 2020), requiring specialized sampling or approximation techniques to maintain computational tractability in real-world applications.

In this paper, we propose a novel graph-based DL model to improve the prediction of demographic and clinical variables from structural brain connectivity. We introduce a computationally efficient and robust architecture that combines three key components: 1) GCNs with residual connections (ResGCN (Pei et al., 2022)) to support brain graph data processing and analysis, 2) fully connected (FC) layers followed by non-linearity for efficient structure-independent feature learning, and 3) a novel Connectivity Attention Block (CAB) that dynamically prioritizes informative inter-regional brain connections. Attention is widely used in recent methods, is competitive with convolutions (Bahdanau, 2014; Vaswani, 2017), and is used in brain image analysis (Jiao et al., 2023; Ranjbarzadeh et al., 2021; Y. Zhang, Teng, et al., 2023). Therefore, the design involving attention allows the model to capture rich representations at both the node (brain region) and subject levels, bridging fine-grained neuroanatomical patterns and global connectivity profiles. Following our previous work demonstrating the improvement that structural connectivity data brings to cortical registration (Zhou et al., 2026) and segmentation (L abiak et al., 2026), here we show how it can help in prediction. Experimental results on two publicly available neuroimaging datasets show that our method outperforms both traditional ML approaches and state-of-the-art graph-based DL models. A preliminary conference version of the proposed network architecture (excluding the new CAB) for the task of sex classification has been previously presented (Kazi, Mora, et al., 2023).

The architecture introduces a combination of complementary components that collectively overcome these individual weaknesses of traditional methods. The integration of ResGCNs with residual connections (Pei et al., 2022) is expected to tackle the over-smoothing problem that plagues standard GCNs in deep architectures, enabling the model to retain discriminative power across multiple layers. The inclusion of FC layers provides structure-independent feature learning capabilities, compensating for the limited generalization issues inherent in graph Laplacian-dependent methods by allowing the model to capture global patterns beyond local graph topology. Most importantly, the novel CAB represents a departure from traditional uniform message aggregation to adaptive, data-driven connection weighting, directly addressing the challenge of dense brain connectivity networks where standard GNN methods exhibit performance degradation. This attention-based approach aligns with recent advances in topological cycle graph attention networks that delineate functional backbones within brain functional graphs by distinguishing key pathways essential for signal transmission from non-essential, redundant connections. By combining these three components, the architecture creates a robust framework that maintains computational efficiency while capturing both fine-grained neuroanatomical patterns and global connectivity profiles, effectively bridging the gap between the theoretical capabilities of GNNs and the practical demands of connectomic analysis for age and cognitive-assessment prediction.

The remainder of this paper, which provides a comprehensive evaluation of the proposed architecture and its implications for neuroimaging research, is organized as follows. We begin with a detailed pre-sentation of the model architecture in the Method section, including the mathematical formulations of all the model components, the design specifications of the FC layers, and the new CAB mechanism. Subsequently, we describe the experimental design in the Experiments section, encompassing dataset descriptions, preprocessing protocols, evaluation metrics, implementation details, and baseline comparisons with both traditional ML approaches and state-of-the-art graph-based DL models for cross-validation and test data. The Results and Discussion section presents quantitative performance analyses across multiple neuroimaging tasks, including age and MMSE score prediction, along with ablation studies demonstrating the contribution of each architectural component. Following the empirical evaluation, we provide a thorough discussion of the findings, addressing the model’s strengths, limitations, and computational efficiency compared to existing methods. Finally, we explore the clinical implications of our approach, discussing how the improved prediction accuracy and interpretability features of the attention-based connectivity analysis can inform clinical decision-making processes and contribute to a better understanding of brain aging and patterns of cognitive decline in neurological and psychiatric disorders.

## 3 Method

Given a dataset of *S* subjects, each subject has an associated brain graph **G** ∈ ℝ*^N^*^×^*^N^* with *N* denoting the number of brain regions (graph nodes), a feature matrix **X** ∈ ℝ*^N^*^×^*^M^* with *M* the number of features per node, and a label *y* ∈ ℝ (i.e., value of the clinical or demographic variable to be predicted). The task is to predict *y*, for which we define a model *f_θ_*as:

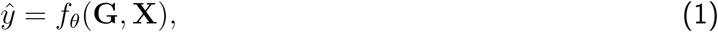

where *θ* is the set of learnable parameters. The model we propose comprises four sub-parts, modules 1 to 4, as shown in Figure 1. Each module is designed to process the combination of **X** and **G** to produce a latent embedding. The last part combines all the previous outputs to produce the predicted label *y*^. We describe all four modules separately and then explain the whole end-to-end model.

**Figure 1:**
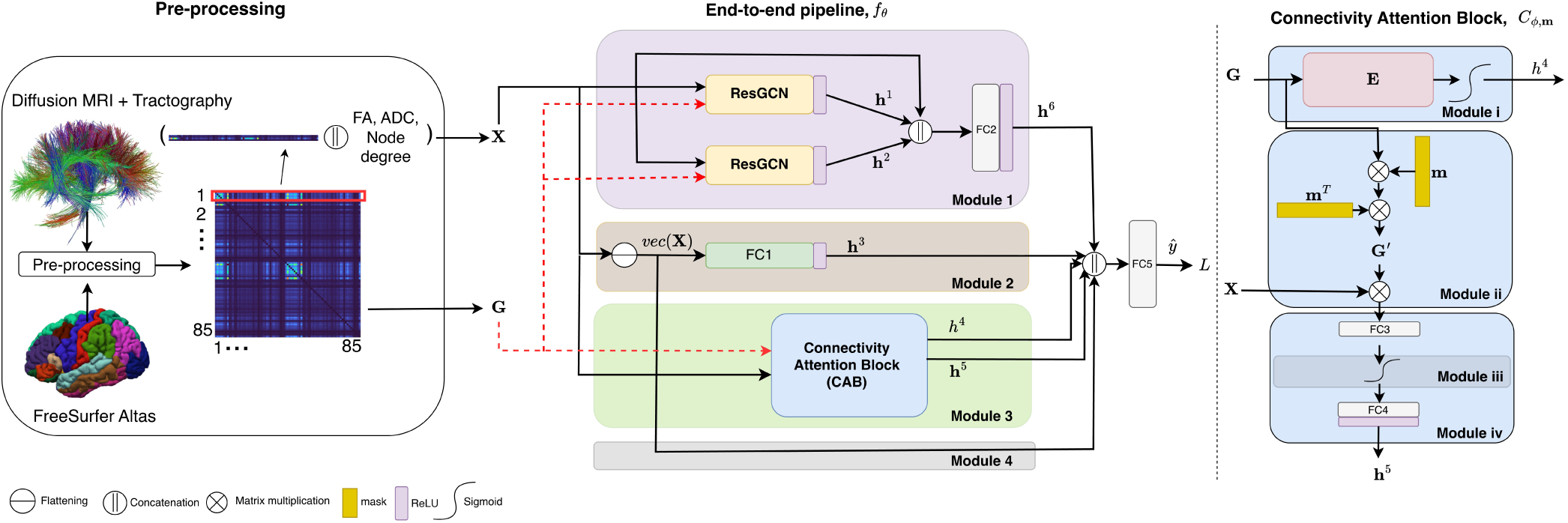
Overview of an end-to-end DL architecture for brain connectivity analysis using dMRI and tractography. The pre-processing module (left) includes the dMRI-based tracts forming the structural connectome matrices. dMRI-derived features (FA, ADC, degree) are concatenated with each row of the connectivity matrix and then used as node features. The model (middle) processes input features and graphs through residual GCNs, fully connected (FC) layers, and a novel Connectivity Attention Block (CAB, right), which refines the graph structure and highlights key connections. Outputs from all modules are combined to predict phenotypes. See the Method section for more details.

### 3.1 Module 1: Graph Convolution

This module comprises two branches, each with a graph convolution with a different embedding size, and a skip connection as shown in Figure 1. We employ a combination of ResGCN layers (Bresson & Laurent, 2017), which captures low-level features of the graph. Each ResGCN (with a different embedding size) transforms the same input data into a different space, enabling the model to learn diverse representations of the graph data. One layer focuses on capturing low-dimensional, essential features, while the other learns higher-dimensional, more nuanced relationships. The skip connection is then combined with these learned representations, allowing the model to leverage both the basic, essential features and the more intricate relationships within the graph. The outputs of these layers are fed to an FC layer (shown as FC2 in Figure 1) to produce the output of the module, **h^6^**.

### 3.2 Module 2: Fully Connected (FC1) Layer

In this layer, we remove any structure that is present in the feature vector sequence **X** and simply extract information from the raw data. This layer can be mathematically defined as: **h^3^** = *ρ* (*FC*1(*vec*(**X**))), where *vec*(**X**) ∈ ℝ*^NM^*^×1^ is the flattened matrix **X**, and *ρ* is a non-linearity (ReLU). An FC layer processing node in parallel with GCN layers offers several potential benefits. It allows the model to learn an independent representation of the node features, capturing information that might not be directly reflected in the graph structure. This is particularly advantageous when the node features themselves hold significant information for the task at hand.

### 3.3 Module 3: Connectivity Attention Block (CAB)

We propose a customized CAB, *C_ϕ,_***_m_**, so that the model learns an embedding representation of a brain graph, as well as to provide a graph-level attention mechanism. The CAB, depicted in Figure 1 (right), is represented as

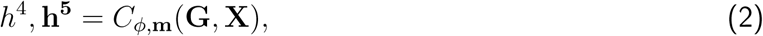

where *h*^4^ is a scalar embedding of each subject’s brain connectivity, which is also treated as a factor of importance of each subject with respect to the population. **h^5^** is a lower-dimensional representation that is sculpted out of **G** and **X**. The external trainable parameter **m**, initialized uniformly, is the attention mask assigned to the nodes (i.e. regions). All are defined as:

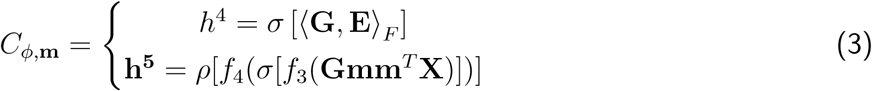

*h*^4^ is the Frobenius inner product of the **G** matrix by a weight matrix **E** ∈ *ϕ* learned by the model, followed by sigmoid non-linearity, *σ*. For **h^5^**, we effectively project **G** onto a single one-dimensional orientation within the *N*-dimensional space, from which we create a rank-one matrix **G**^′^: = **Gmm***^T^*, with the projection weights **m** learned by the model. *f*_3_ and *f*_4_ are the FC layers *FC*3 and *FC*4 as shown in Figure 1, both followed by non-linearities. Here, instead of applying attention to each element of **G**, we leverage the matrix properties of **G**. Our goal is to retain a single representative rank-one matrix **G**^′^ that best helps the prediction task at hand. This operation simplifies the representation of the matrix while preserving its most significant features, along the same lines as singular-value decomposition. The vector **m** is optimized jointly with the rest of the network to maximize prediction performance, by weighting the region-wise features by **m** and projecting the results onto the direction **Gm** (i.e., eigenvector of **G**^′^), effectively narrowing the attention to a single orientation in the *N*-dimensional space.

Importantly, CAB is not intended to serve as the sole source of model expressivity. Higher-order graph structure and complex interaction patterns are captured by the residual GCN blocks and the fully connected branches (Modules 1 and 2), while CAB provides a complementary, global, and interpretable reweighting of brain regions and their associated connections. This design allows CAB to enhance the model’s representational capacity without replacing the richer structural modeling performed by the other components.

CAB works directly on the full adjacency matrix *G* and learns a single dominant eigen-direction-like *m* (a region mask) to produce a task-adaptive rank-1 approximation *G*^-^ = *Gmm^T^*, distilling global structural connectivity into a low-dimensional, interpretable embedding. This allows CAB to capture global patterns that local/node-level attentions cannot. To compare with existing methods: GAT (Veličkovíc et al., 2017) computes attention only between neighboring nodes (local 1-hop edges), whereas CAB derives the attention weights from the entire connectivity matrix. Low-rank attention (S. Wang et al., 2020) methods approximate sequence attention via query-key projections. CAB instead leverages the symmetry of brain connectivity matrices via a rank-1 projection, avoiding unnecessary decompositions. Spectral attention (N. Liu et al., 2024) methods rely on fixed Laplacian filters. CAB learns a data-driven eigen-like vector end-to-end, optimized directly for the prediction task.

### 3.4 Module 4: Skip Connection

We add an overall skip connection (Module 4 in Figure 1) to help mitigate the vanishing gradient problem and allow effective information propagation across layers by directly connecting *vec*(**X**) to the final layer.

### 3.5 Data Fusion and Loss Function

The outputs of all four modules are concatenated as **X_all_** = [**h^3^***, h*^4^, **h^5^**, **h^6^***, vec*(**X**)] and fed to an FC layer to produce the final prediction *y*^ = *FC*5(**X_all_**)

For the prediction task, we use a customized loss function made of two parts. The first part is the Huber loss, *L_δ_*(*y* − *y*^), which is a piecewise continuous function, defined as:

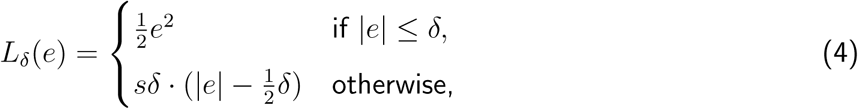

where *δ >* 0 is a threshold that controls the location of the transition between quadratic and linear loss behaviors. This function applies a quadratic loss when the absolute error is less than or equal to *δ* and a linear loss when the error exceeds this threshold, effectively combining the differentiability of the quadratic loss at small errors with the robustness of the linear loss to outliers. The second term of the loss, *L^R^*, is the regularizer for **m** defined as *L^R^ = Σ^N^_i_ | m_i_| Σ^N^_i_ | m_i_| log | m_i_|* (Kazi, Farghadani, et al., 2023). The overall loss for the model is therefore L = Lδ + βL^R^, where β > 0 is a weighting factor.

## 4 Experiments

### 4.1 Datasets

We use two publicly available datasets. The details are given below.

**Pre-symptomatic Evaluation of Experimental or Novel Treatments for Alzheimer’s Disease (PREVENT-AD)** (Leoutsakos et al., 2016) is a publicly available dataset that provides a comprehensive set of data on individuals who are at risk for developing AD (https://prevent-alzheimer.net). The database contains neuroimaging studies such as MRI (including dMRI) and PET scans, a range of demographic, clinical, cognitive, and genetic data, as well as data on lifestyle factors such as diet and exercise. The dataset comprises 347 subjects, some with multiple (longitudinal) dMRI scans, totaling 789 dMRI scans.

The third release of **Open Access Series of Imaging Studies (OASIS3)** (LaMontagne et al., 2019) is a longitudinal neuroimaging dataset with clinical and cognitive scores for normal aging and AD, and is provided freely to researchers (http://www.oasis-brains.org). The OASIS3 dataset contains MRI scans (including dMRI), cognitive assessments, demographic information, and clinical diagnoses for subjects, including healthy controls, individuals with mild cognitive impairment, and AD patients. From this dataset we used 1294 brain scans from 771 subjects. Please refer to Table 1 for the overview of the datasets.

**Table 1:**
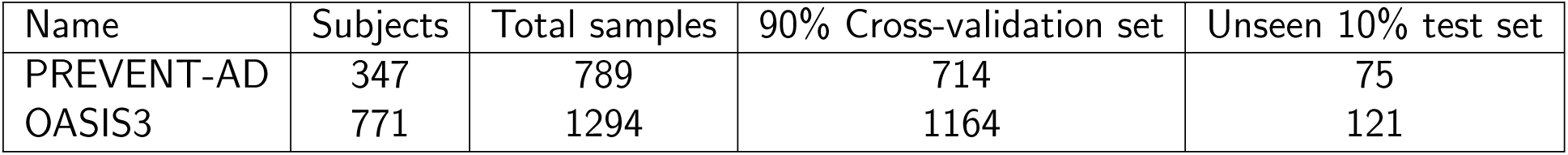
Description of dataset sizes (number of available scans) and partitioning for PREVENT-AD and OASIS3 dataset. Due to missing demographic data, nine subjects were removed from the OASIS3 dataset. Each subject may have multiple samples collected at different time points, hence there are overall more samples than subjects.

#### 4.1.1 Pre-processing

We used FreeSurfer (Fischl, 2012) to process the databases (additionally applying the longitudinal processing pipeline (Reuter et al., 2012) for PREVENT-AD). We then ran the FreeSurfer diffusion processing pipeline and propagated the 85 automatically segmented cortical and subcortical regions from the structural to the diffusion space. These 85 regions act as the nodes in our graph setup. Next, we used the public CSA-ODF and Hough Tractography toolbox (http://www.nitrc.org/projects/csaodf-hough) to reconstruct the diffusion orientation distribution function in constant solid angle (Aganj et al., 2010), run Hough-transform global probabilistic tractography (Aganj et al., 2011) to generate 10,000 optimal streamlines per subject, compute symmetric structural connectivity matrices, and augment the matrices with indirect connections (Aganj et al., 2014). More details on the pipeline can be found in our previous publication (Aganj et al., 2023). Once we had all the graphs *G_i_*, we performed a population-level normalization on edge weights.

For node features, we used the structural connectivity map of the node region to the rest of the regions as well as mean fractional anisotropy (FA), mean apparent diffusion coefficient (ADC), volume, and the degree of the node. In predicting age and dementia, dMRI analysis can benefit from a combination of node features. FA and ADC, reflecting white matter organization and water diffusion, respectively, capture microstructural changes due to aging and AD, whereas mean segmentation volume captures brain region size that is potentially reduced in age-related atrophy or AD. Finally, the node degree (total connectivity to a brain region) reveals alterations in brain network connectivity patterns. Therefore, for each subject, we obtained **G***_i_* ∈ ℝ^85×85^ and corresponding **X***_i_* ∈ ℝ^85×89^ (i.e., *N* = 85 and *M* = 89). The four additional features capture complementary information beyond connectivity and standard diffusion measures, improving robustness and performance. FA captures the degree of directional water diffusion and serves as a microstructural marker of white-matter organization, whereas ADC reflects the overall magnitude of diffusion within tissue. The regional volume feature quantifies morphometric variation across the segmented brain regions, and node degree represents the number of structural connections incident to each node, thereby summarizing its topological centrality within the connectome.

#### 4.1.2 Implementation Details

All the experiments were run via 10-fold cross-validation with the same folds across methods and experiments. For model robustness, we added zero-mean Gaussian noise with a standard deviation of 0.1 to the training samples. All the experiments were run on a Linux machine with 512 GB of RAM, an Intel (R) Xeon (R) Gold 6256 CPU @ 3.60 GHz, and an NVIDIA RTX A6000 (48 GB) graphics processing unit. The total number of parameters used in the proposed model was 5073, which was comparable to GCNConv (2453), DGCNN (4653), GraphConv (4653), ResGatedGraphConv (RGGC) (9128), and GINConv (2683). In our experiments, the output embedding sizes for **h^1^**, **h^2^**, **h^3^**, and **h^5^** were 25, 20, 5, and 2, respectively. The values of *β* = 0.01 chosen empirically in the loss function. Each input feature and edge weight was normalized using a fixed population-level min–max scaling, where the range was computed from the combined training and validation (but not the test) cohorts. This ensures a consistent scale across features and prevents high-variance attributes (e.g., volumetric measures) from disproportionately influencing the model gradients.

### 4.2 Experimental Design

#### 4.2.1 Evaluation Metrics

Performance is evaluated using four complementary metrics: Root Mean Square Error (RMSE) and Mean Absolute Error (MAE), which measure the magnitude of prediction errors, as well as Pearson’s Correlation Coefficient (PC) and Spearman’s Correlation Coefficient (SC), which assess the strength and direction of the linear and rank-order relationships, respectively, between predicted and ground-truth values. RMSE is particularly sensitive to large errors, making it useful for penalizing significant deviations, while MAE provides a more interpretable average error magnitude. Pearson’s correlation captures linear associations, whereas Spearman’s accounts for monotonic but potentially non-linear trends. Together, these metrics offer a comprehensive assessment of both the accuracy and reliability of the model predictions across all three tasks, enabling a nuanced understanding of performance from both error-based and correlation-based perspectives.

#### 4.2.2 Cross-validation and Unseen Test Set Experiments

We divide the entire dataset into a 90:10 split. We set aside the 10 % data as an unknown test set and only use it once the training of the model is done. We use the remaining 90% data for cross-validation by splitting the data again into 90:10 ratio. The dataset split is shown in Table 1. We use sklearn (Feurer et al., 2020) to perform 10-fold cross-validation using experiments for six classical ML and 5 DL based methods. All the sets are created based on subject IDs instead of sample IDs to avoid data leakage and ensure that no information from the same subject appears in both the training and validation/test sets. We keep 10% of the data aside from each dataset so as not to heuristically fit the model to the entire data, and then test the model at the end on the unseen data, the results of which are reported in the next section.

#### 4.2.3 Baselines and Comparative Methods

We performed a comparative evaluation of various conventional ML and advanced DL models for the age prediction task across the two datasets of PREVENT-AD (789 scans) and OASIS3 (1294 scans), as well as for MMSE prediction in the OASIS3 dataset, while ensuring that all scans of the same subject appeared only in a single (train, validation, or test) set. As a baseline, we employ a naive model that consistently predicts the mean (resp. median) of the training set. We evaluate its performance using RMSE (resp. MAE) on the validation set. These baseline metrics provide a sanity check, establishing the minimum performance threshold that all other models are expected to surpass. Since this model outputs a constant value, its correlation coefficients (PC and SC) with the ground truth are zero. The conventional ML models that we tested include linear regression (LR), support vector regression (SVR), decision trees, regression trees, ensemble trees, and neural networks (all implemented in the sklearn package). We further compared our method with DL approaches such as the multilayer perceptron (MLP), and more advanced GNN models such as GCN, GIN, GraphConv, and ResGCN.

#### 4.2.4 Ablation Studies

We performed ablation experiments to systematically evaluate the individual contributions of each architectural component in the proposed model and to understand their impact on overall predictive performance. By selectively removing key modules—such as the FC1 block, CAB block, skip connection, and GCN block—and observing changes in prediction accuracy, we aimed to identify which components are essential for capturing complex patterns in the data. This approach helps to validate the necessity of each element within the model design, ensuring that the architecture is both effective and interpretable. Moreover, it highlights the role of graph-based learning in enhancing the model’s capacity to model relationships within neuroimaging data, thereby justifying the inclusion of the GCN block in particular. We used the same evaluation metrics as for the comparative methods.

#### 4.2.5 Qualitative Results

We illustrate the scatter plots of the predictions vs. ground-truth data for the three tasks for the test set. We further report the significance of our results using a paired *t*-test between the folds.

## 5 Results and Discussion

### 5.1 Cross-validation and Unseen Test Set Experiments

Table 2 shows the results on the validation set of the 10-fold cross-validation for the age prediction task on the PREVENT-AD and OASIS3 datasets. The table is divided into three parts horizontally. The first part consists of the naive model. The mean RMSE is higher for the OASIS3 age prediction task due to its wider distribution compared to PREVENT-AD. However, the standard deviation of RMSE relative to its mean is smaller in OASIS3, possibly due to its larger dataset size. A larger dataset can provide more stable and reliable estimates of the true metric values, leading to reduced variation across cross-validation folds. The next set showcases the performance of 6 traditional ML methods. For the PREVENT-AD dataset, we observe that only the Ensemble Tree is able to outperform the naive baseline model. Where as for the OASIS3 dataset, most of the traditional ML methods perform better than the naive classifier. The third part of the table is (mostly graph-based) DL methods. Most of the methods in this category show better performance than the naive model across both datasets. MLP, being the only non-graph DL method, performs better than almost all of the traditional ML methods. The MLP’s layered structure allows it to hierarchically extract and combine features, making it more expressive and better suited for modeling non-linear aging trajectories and interdependent biomarkers. Most of the graph DL methods perform better than traditional ML methods and MLP. This improved performance can be attributed to the ability of GNNs to model relational structures and population-level dependencies that are otherwise ignored in traditional or naive models. GNN-based methods incorporate subject-level connectivity information and leverage inter-subject similarities through graph representations. These methods can effectively capture non-linear interactions and high-dimensional spatial patterns in the data—especially useful in aging neuroimaging datasets, as brain connectivity is affected in aging. Furthermore, the inductive bias introduced by graph convolutions allows the models to generalize better to unseen subjects, particularly when brain regions or connectivity features exhibit complex dependencies. This capability likely explains their superior correlation with actual age, even when RMSE values are only moderately improved compared to simpler models. The proposed model consistently demonstrated superior performance, validating the strength of the architecture in capturing graph-structured population-level information across heterogeneous datasets. The proposed model integrates the strengths of MLPs (non-linear feature learning), ResGCNs (deep hierarchical graph representation), and a CAB (adaptive weighting of informative connections), enabling it to outperform all baseline methods by effectively capturing both individual and population-level patterns. The asterisk symbol shows where the proposed model performs significantly better based on the paired *t*-test (*α* = 0.05).

**Table 2:**
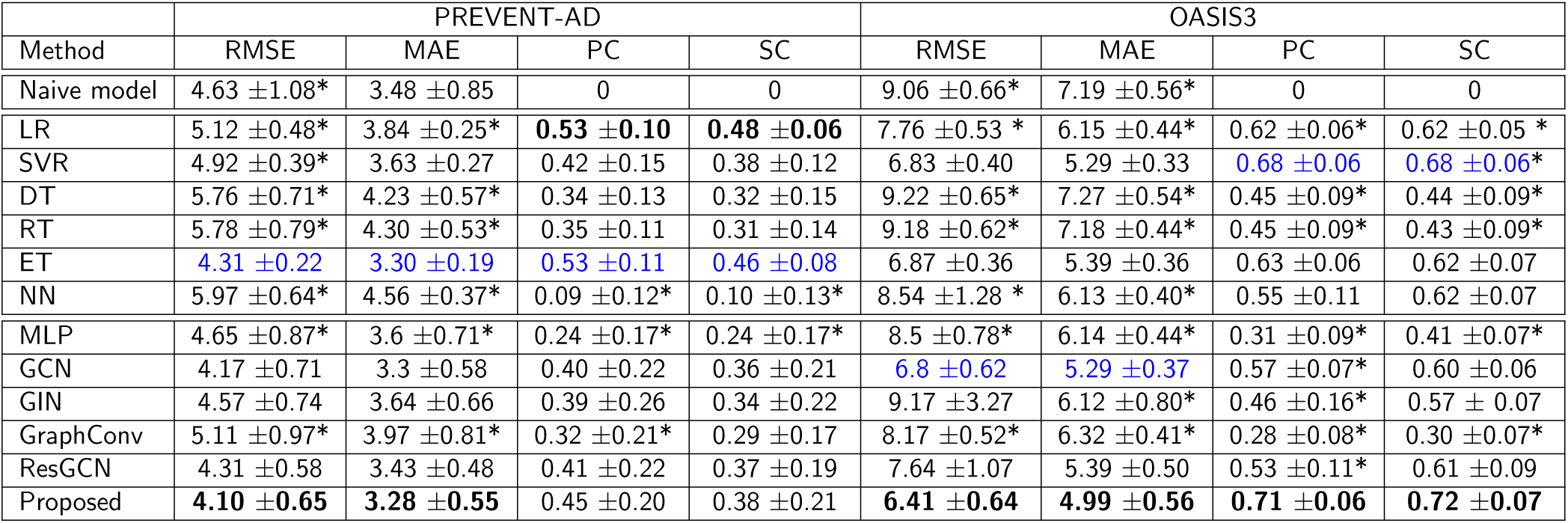
Age prediction results on the validation set from the 10-fold cross-validation, for the PREVENT-AD and OASIS3 datasets. Asterisk ‘*’ indicates that the proposed method significantly outperformed the respective methods when compared using paired *t*-test (with *α* = 0.05). Bold and blue indicate the **best** and second-best, respectively, in each column. LR = Linear Regression (Su et al., 2012); Support Vector Regression (SVR) (Awad & Khanna, 2015); DT = Decision Tree (De Ville, 2013); RT = Regression Tree (Loh, 2011); ET = Ensemble Tree (Che et al., 2011); NN = Neural Network (S.-C. Wang, 2003); MLP (Goodfellow et al., 2016); GCN (Kipf & Welling, 2016); GIN (K. Xu et al., 2018); GraphConv (Fey & Lenssen, 2019); ResGCN (Pei et al., 2022).

Similarly, Table 3 shows results on MMSE prediction on OASIS3 dataset. We can see a trend similar to Table 2, except that SVR performs best and the proposed method is mostly second best. Most of the traditional ML methods perform worse than the baseline, whereas the performance metrics of the DL methods are marginally better. This can be explained with a histogram of the ground-truth values, plotted for each dataset and task in the Figure. 2. As can be seen, MMSE has a much more skewed distribution than age does, potentially explaining the poor performance for the MMSE task. Although GNNs are effective for modeling complex relational patterns, MMSE scores are sparse, bounded, and heavily concentrated at higher values in healthy subjects, resulting in a skewed distribution and limited relational signal. For this type of endpoint, conventional models such as SVR and ensemble tree methods were able to capture the subtle variations adequately, and the added complexity of GNNs provided little additional benefit.

**Figure 2:**
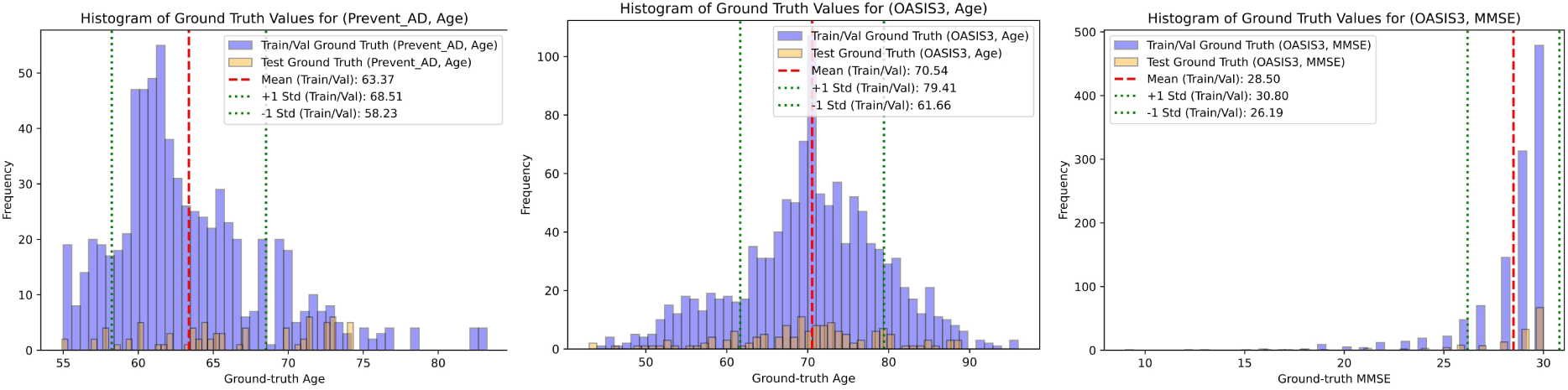
Histograms of the ground-truth values in the cross-validation (purple bars) and test (orange bars) sets for all datasets and tasks. The mean of each cross-validation distribution is marked by thick red dotted lines, while the standard deviation is indicated by thin green dotted lines. These visualizations help assess how well the training and test distributions align and provide insights into potential challenges in model training. For the PREVENT-AD dataset, the age distribution is notably skewed, indicating an imbalance in subject age representation. The OASIS3 dataset’s MMSE distribution exhibits a power-law trend, where most samples cluster in the higher score range (i.e., cognitively normal individuals), with relatively few samples showing lower scores. The age distribution in the OASIS3 dataset is approximately symmetric and Gaussian, which is more favorable for training.

**Table 3:**
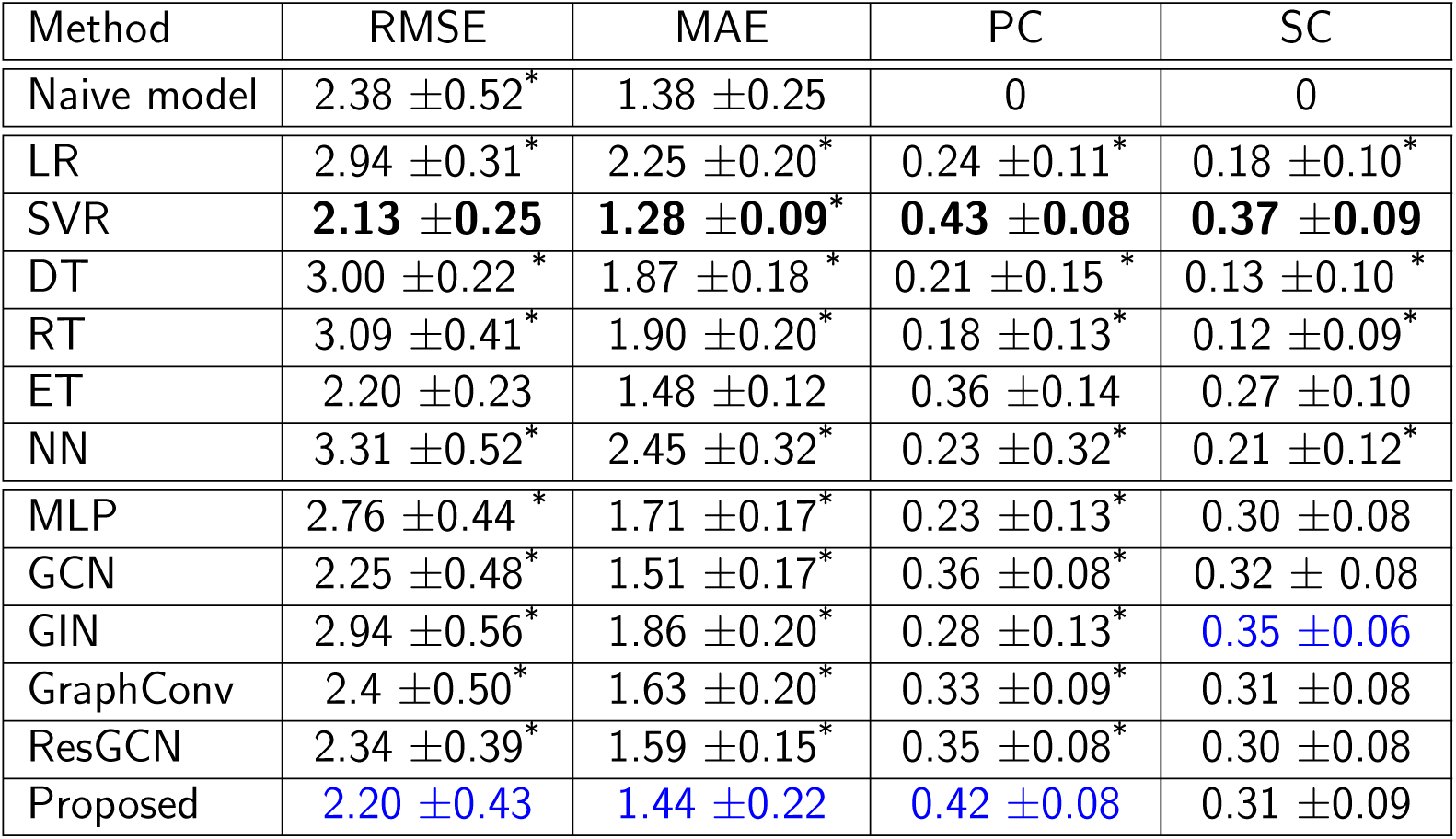
Cross-validation results by all methods for the MMSE prediction task on the validation set using the OASIS3 dataset. Bold and Blue indicate the **best** and second-best, respectively, in each column. For the fullforms and corresponding citation please refer to Table 2.

### 5.2 Results on 10% Unseen Test Data

We hypertune all the models during cross-validation, and then use the best setup for each model to evaluate its performance on the unseen test dataset. In these (single-fold) experiments, we train the model on all the cross-validation data and test it on the held-out test data. For each fold in the cross-validation, we stop the training at the epoch yielding the lowest loss on the validation set. For the held-out test data, we stop the training at the same epoch as that of the model corresponding to the lowest loss among the folds. For both datasets, as shown in Table 4, the proposed method outperformed all other models in age prediction in 7 out of 8 metrics, achieving the lowest prediction error (except it finished second in MAE for PREVENT-AD) while also demonstrating the highest correlation scores, indicating superior predictive accuracy. For MMSE prediction (in OASIS3), the proposed method still outperformed the other DL methods, but not the conventional ML methods of Ensemble Trees and (mostly) SVR. The narrow range of MMSE in the OASIS3 dataset (as reflected in the Naive model row and Figure 2) resulted in performance metric values that were close to each other among models.

**Table 4:**
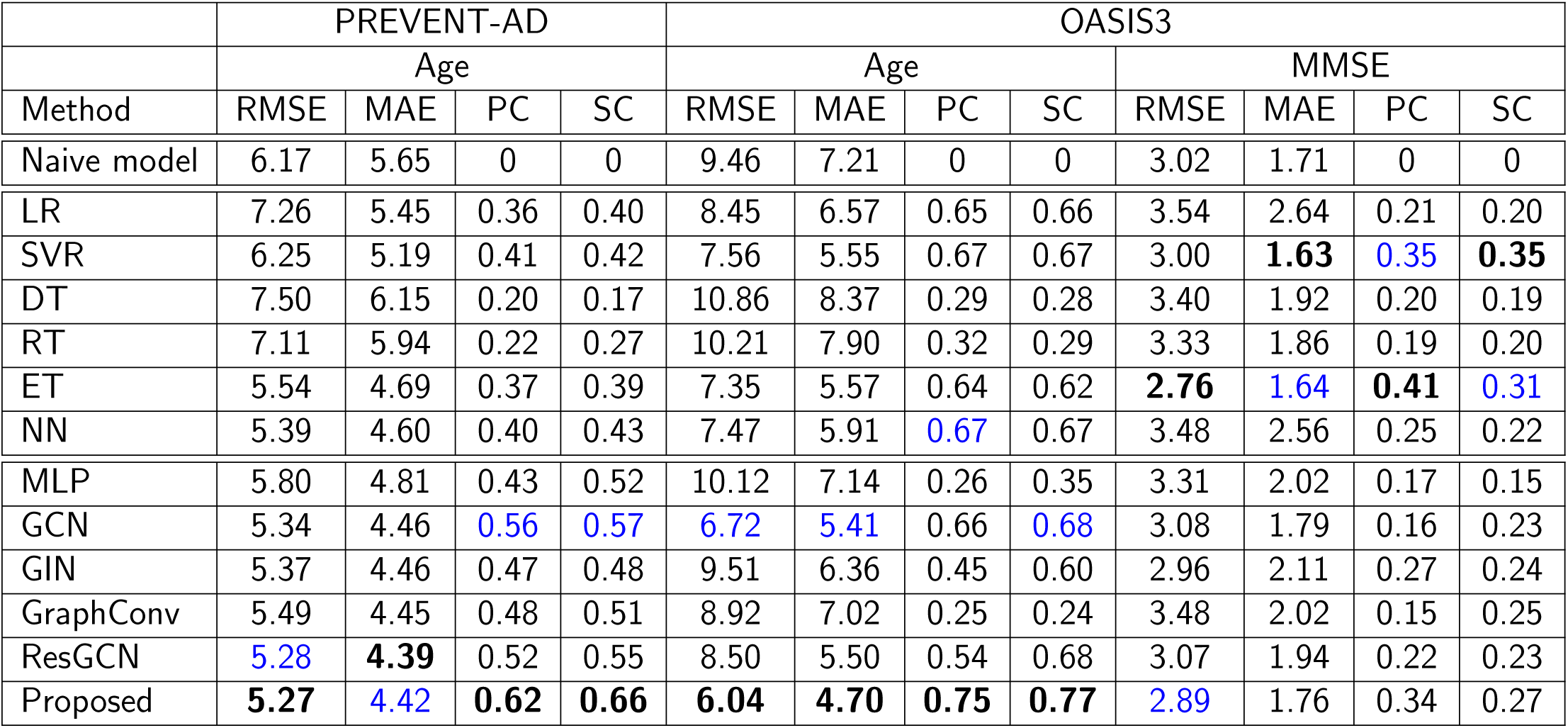
Performance evaluation for all the methods on 10% unseen test data. Bold and Blue indicate the **best** and second-best, respectively, in each column. For the fullforms and corresponding citation please refer to Table 2.

We additionally performed these experiments on data from the second phase of the Alzheimer’s Disease Neuroimaging Initiative, with a dataset size of 200 samples. While the proposed method still outperformed the rest of the methods, its prediction error was not substantially below the baseline, presumably due to the small dataset size (PC/SC were still 0.50/0.55 and 0.34/0.36 for age and MMSE, respectively).

### 5.3 Ablation Tests

Table 5 presents an ablation study on the prediction tasks on both datasets for the test set. For PREVENT-AD, when removing specific components such as the FC1 block, CAB block, skip connection, or GCN block, performance degrades across all datasets, confirming the effectiveness of the model’s complete architecture. The trend remains generally consistent in OASIS3 (age); however, the original model without the FC1 component maintains the best scores, whereas other ablations cause notable declines in performance.

**Table 5:**
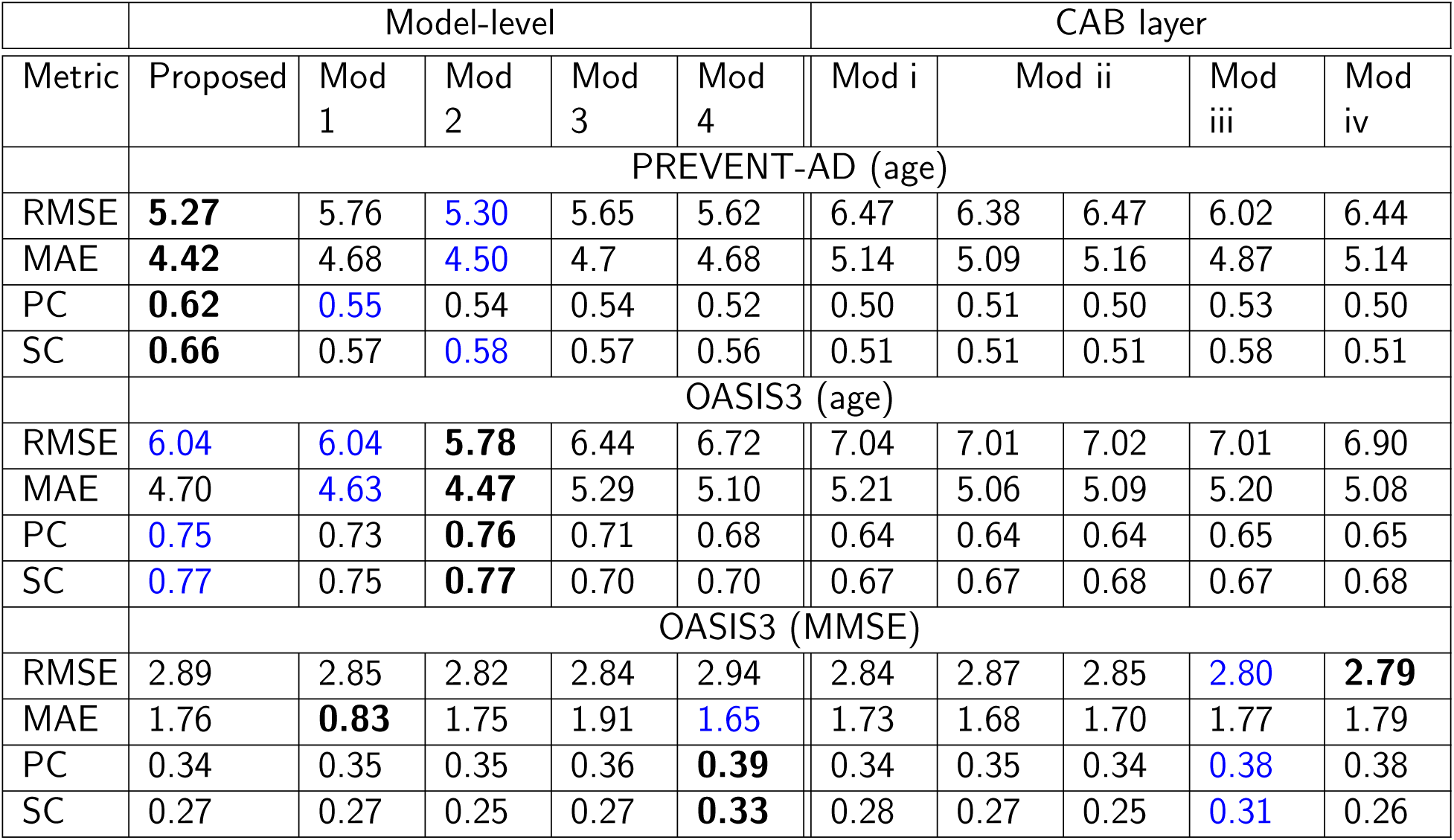
Ablations on the prediction tasks. Each column represents the respective block removed from the pipeline. Bold and Blue indicate the **best** and second-best, respectively, in each row. Mod denotes a module ablation. We define: Mod 1 — no ResGCN layers; Mod 2 — no FC1 layer; Mod 3 — no CAB; Mod 4 — no skip connection; Mod i — no *E* in CAB; Mod ii (left) — no *G***mm***^T^* and Mod ii (right) no *G* in *G***mm***^T^*; Mod iii— no Sigmoid in CAB; Mod iv — no MLPs in CAB.

For OASIS3 (MMSE), the original model without the skip connection outperforms the other ablated versions in terms of correlations, though the performance drop is insubstantial. Removing any component results in only a marginal increase in RMSE and a decrease in correlation metrics, possibly due to the narrow range of MMSE values in OASIS3.

For age prediction, components that model global and relational structure are most critical. In particular, removing the GCN block (Mod 1) led to consistent degradation in correlation (e.g., PC drops from 0.62 to 0.55 in PREVENT-AD and from 0.75 to 0.73 in OASIS3), confirming that capturing inter-regional dependencies is essential for modeling brain aging. Similarly, the CAB module (Mod 3) showed substantial impact (e.g., RMSE increases from 6.04 to 6.44 in OASIS3), indicating that adaptive weighting of connections is important for identifying age-relevant connectivity patterns. This aligns with the biological premise that aging manifests as distributed, network-level changes. In contrast, for MMSE prediction, the contribution of individual modules is less pronounced. Ablation results show minimal variation across configurations (e.g., RMSE varies only from 2.79 to 2.94, and PC from 0.34 to 0.39). This suggests that MMSE prediction is less sensitive to architectural choices, likely due to the compressed and skewed distribution of MMSE scores (as shown in Figure 2), which limits the model’s ability to leverage complex representations. Notably, removing the skip connection (Mod 4) slightly improves correlation (PC: 0.34 → 0.39), indicating that simpler representations may suffice for this task.

The ablation results generally emphasize that each architectural component plays a crucial role in maintaining predictive accuracy, with graph-based learning contributing significantly to overall model performance. The GCN block, in particular, appears essential for capturing complex dependencies, as its removal leads to a decline in the age prediction accuracy.

### 5.4 Qualitative Results

We show scatter plots of ground truth vs predictions on the held-out test set for all the datasets and tasks for qualitative evaluation of the performance. Figures 3, 4 and 5 show the scatter plots for age prediction in OASIS3 and PREVENT-AD and for MMSE prediction in OASIS3, respectively. The first two rows show the results by the DL methods, and the last two rows show those by traditional ML methods. The proposed method (top-left) shows the tightest clustering along the identity (45°) line, reflecting the highest correlation and lowest prediction error among all methods. These plots are consistent with the summary measures reported in the tables. A consistent trend among all methods is that the slope of the least-squares line is less than 45°, suggesting a compromise between the mean-predicting behavior of the naive model and ideal prediction.

**Figure 3:**
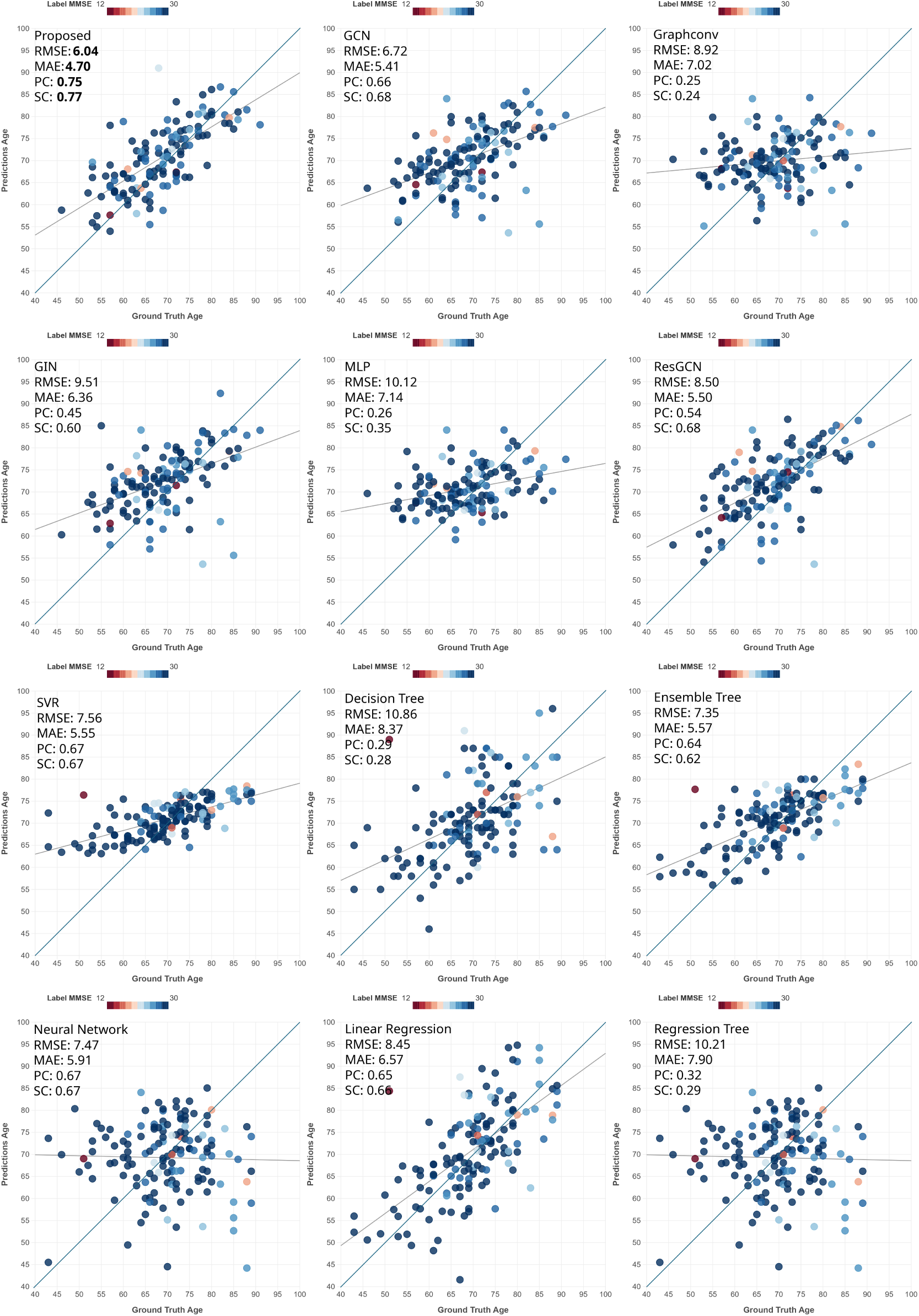
Scatter plots comparing predicted age to ground-truth age on the OASIS3 dataset. Each dot represents a subject in the test set. The darker line is the identity line, whereas the lighter line is the least-squares line. The color indicates the MMSE score.

**Figure 4:**
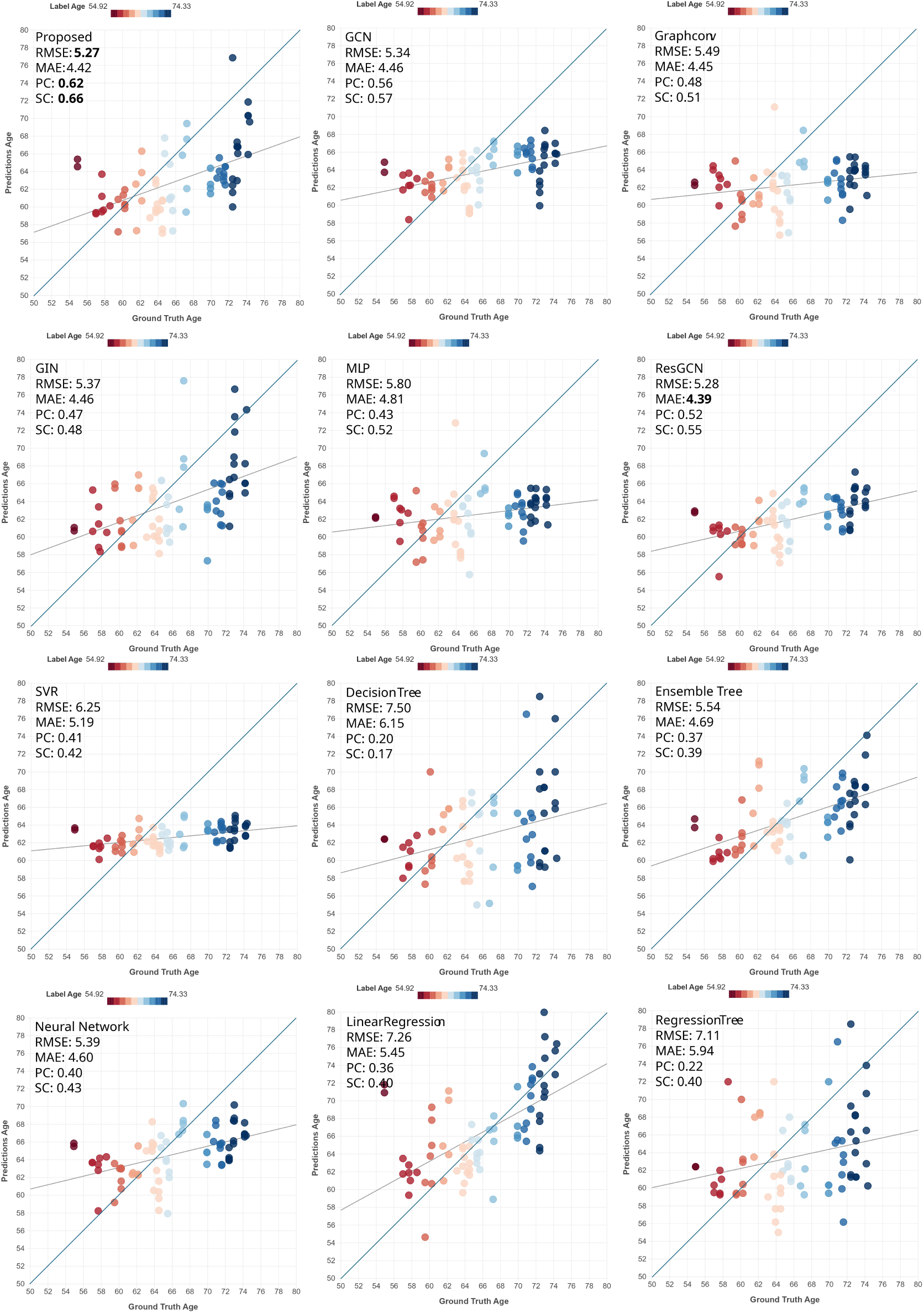
Scatter plots of prediction performance for the test set of the PREVENT-AD dataset for the task of age prediction. The darker line is the identity line, whereas the lighter line is the least-squares line. The color indicates the ground-truth age.

**Figure 5:**
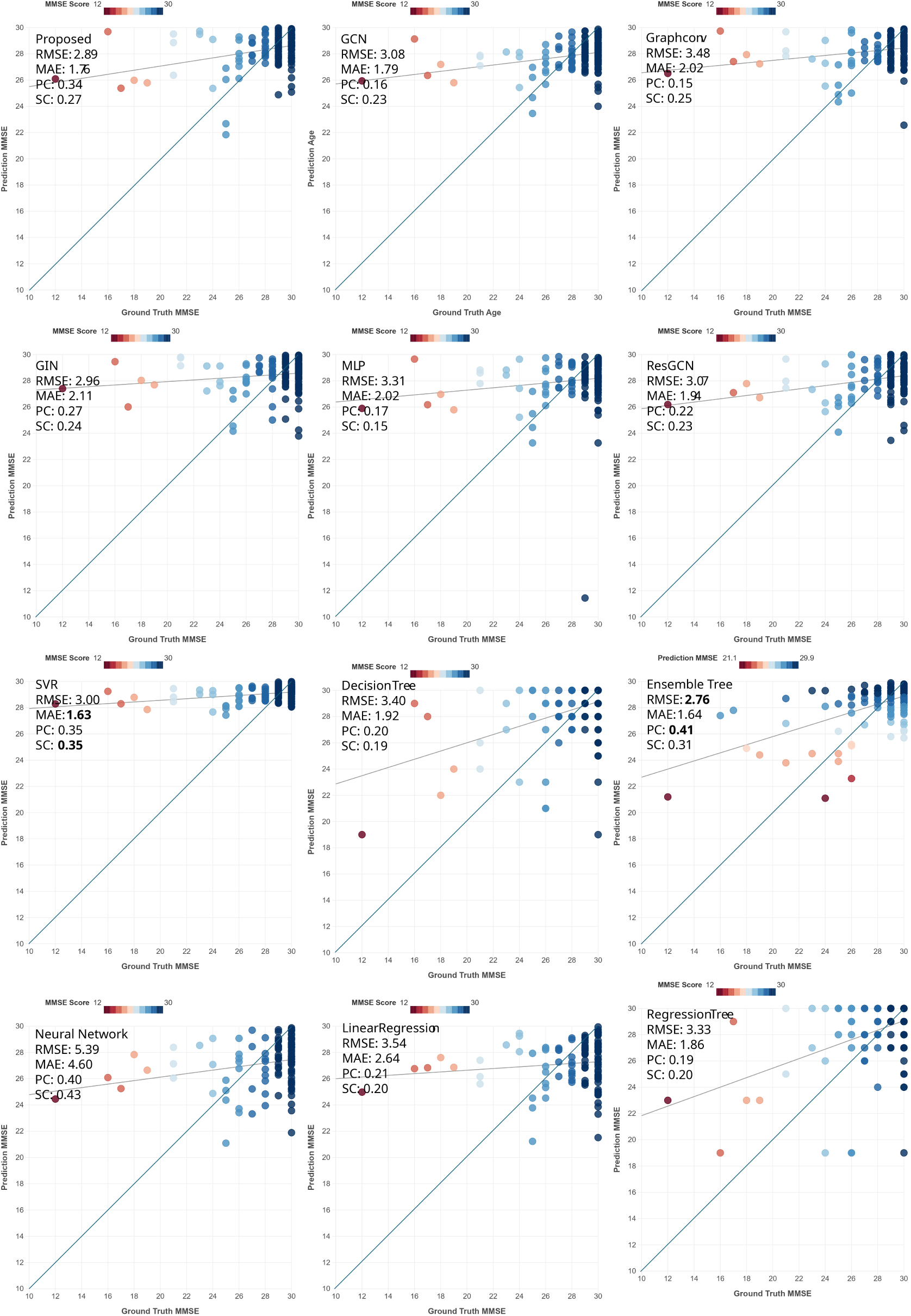
Scatter plots for test set on OASIS3 dataset for the task of MMSE prediction. The darker line is the identity line, whereas the lighter line is the least-squares line. The color indicates the ground-truth MMSE.

### 5.5 Attention learned by CAB

We show the top five regions selected by the CAB in Table 6. For the age task for both datasets, the regions selected and attention values obtained are similar (but not identical) and are different from the MMSE task. As can be seen, the right hippocampus has the strongest weight for age in both datasets.

**Table 6:**
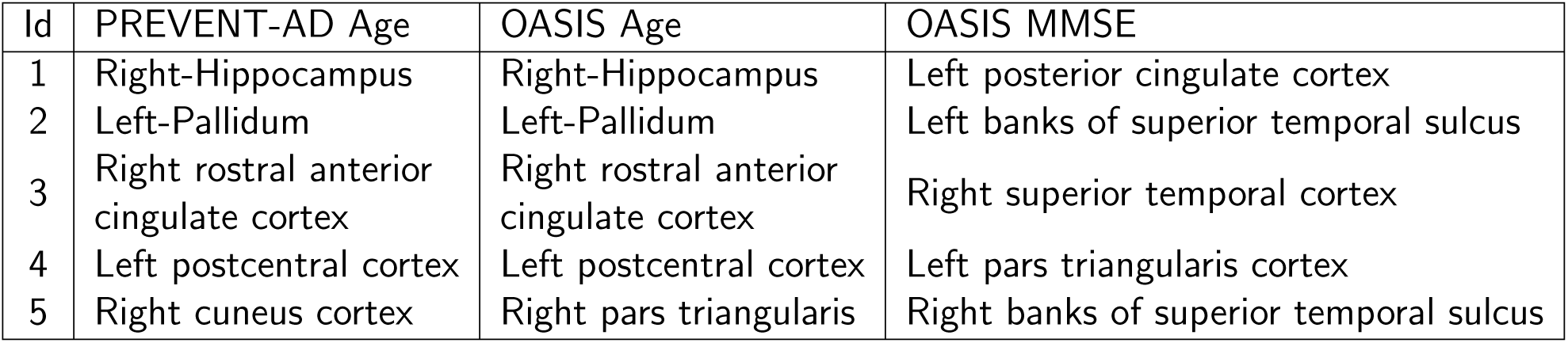
Top regions identified across different tasks.

Literature suggests that the hippocampus is indeed representative of aging (Aganj et al., 2023; Y. Xu et al., 2008); as we have seen in three datasets with aging populations, age was most strongly correlated with hippocampal connectivity (Aganj et al., 2023). Posterior cingulate cortex received the highest weighting for the MMSE task.

### 5.6 Hyperparameter and training time comparison

The output embedding sizes for the five modules were selected empirically through systematic hyperparameter sweeps during initial model development and benchmarking. The row ‘Embedding size’ in Table 7 shows the embedding sizes for the proposed and all the comparative methods.

**Table 7:**
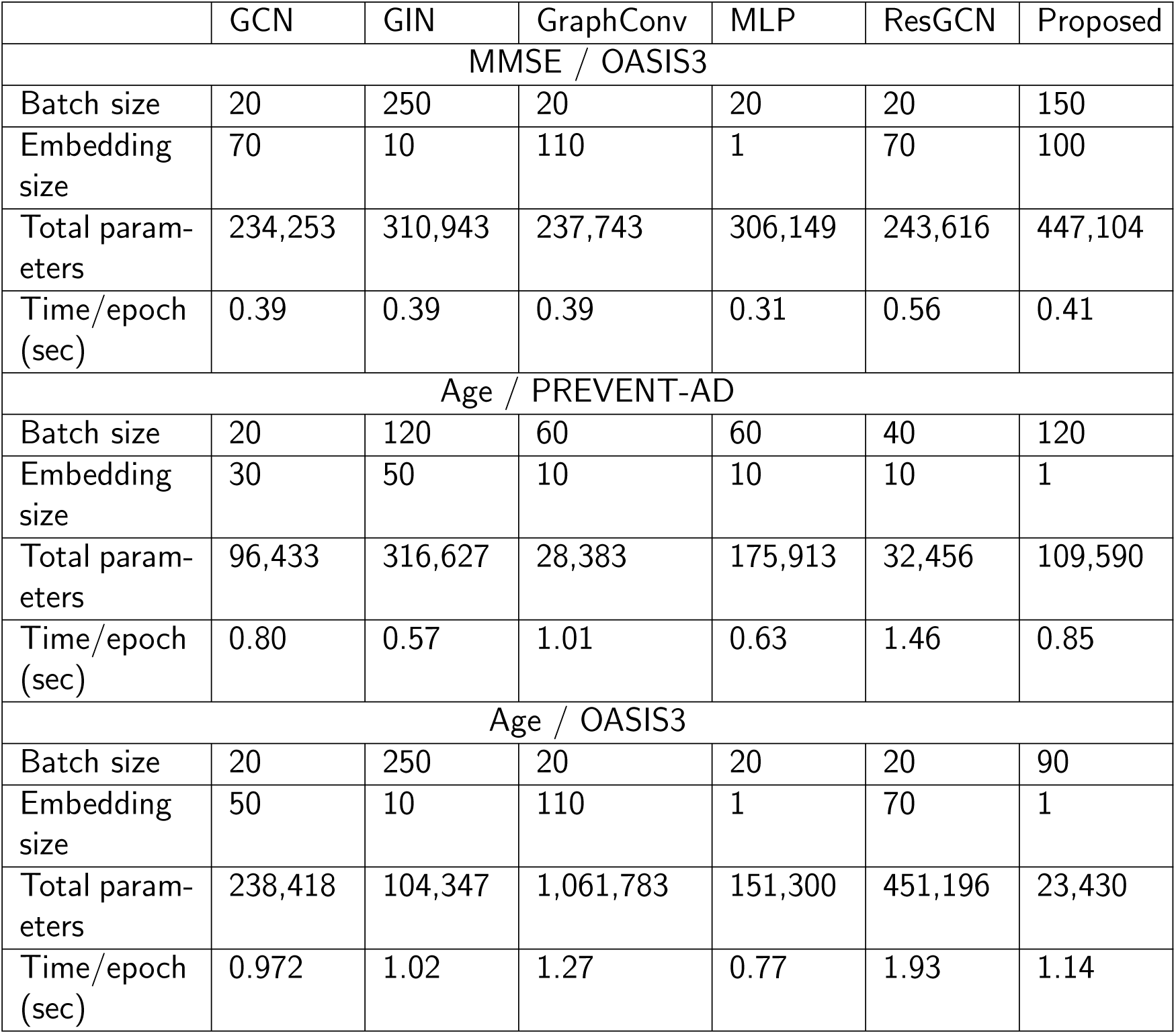
This table summarizes the selected hyperparameters and training efficiency across three setups. The learning rate was set to 0.0001 for all experiments, except for the age task on the OASIS3 dataset, where it was empirically set to 0.001. The proposed model is competitive in runtime, but its parameter count and speed vary by dataset, showing that performance depends strongly on the task.

As shown in Table 7 for the MMSE prediction task on OASIS3, most graph-based models maintain similar training times per epoch despite differences in parameter counts, while the proposed model shows a moderate increase in parameters with only a slight impact on runtime. In the Age prediction task on PREVENT-AD, the proposed model achieves a balanced trade-off, maintaining competitive training time with a moderate number of parameters compared to both lightweight (GraphConv) and heavier (GIN) architectures. For Age prediction on OASIS3, a larger variation is observed, where models such as GraphConv exhibit significantly higher parameter counts, whereas the proposed model remains relatively compact with competitive runtime. Overall, these results highlight that model complexity and computational cost are highly dataset- and task-dependent, and the proposed approach achieves a favorable balance between efficiency and capacity across different experimental settings. The proposed method is robust because it stays competitive in training time while using much fewer parameters in Age/OASIS3 than the other models. It is also consistently near the faster end of the table, which suggests a good balance between efficiency and model capacity.

## 6 Conclusion

In this paper, we proposed a simple yet effective multi-branch model capable of capturing complementary information from structural brain connectivity, which we evaluated in the context of age and MMSE prediction. The configuration of input data and multiple operations from all four modules helped to learn better representations of each subject’s graph. We have shown that our model often outperforms competing techniques on two publicly available datasets, while also ablating all the components of the model. Future work includes: the addition of interpretability to the models to find the brain subnetworks that are informative for the prediction task, as well as trying other (such as gated attention) graph convolution mechanisms; investigation of the suitability of as a subject-level attention score learned by the network, with the goal of quantifying how representative each subject is within the overall population; systematic feature ablation studies across all methods to assess the contribution of each input modality to model performance; robustness analysis of our approach to domain shift and its generalizability by testing it on other clinical datasets after connectivity harmonization (Zhou et al., 2025).

## 7 Acknowledgments

Support for this research was provided by the National Institutes of Health (NIH), specifically the National Institute on Aging (RF1AG068261, R01AG068261).

Additional support was provided in part by the BRAIN Initiative Cell Census Network grant U01MH117023, the National Institute for Biomedical Imaging and Bioengineering (P41EB015896, R01EB023281, R01EB006758, R21EB018907, R01EB019956, P41EB030006), the National Institute on Aging (R56AG064027, R01AG064027, R01AG008122, R01AG016495, R01AG070988), the National Institute of Mental Health (R01MH121885, RF1MH123195), the National Institute for Neurological Disorders and Stroke (R01NS0525851, R21NS072652, R01NS070963, R01NS083534, U01NS086625, U24NS10059103, R01NS105820), the NIH Blueprint for Neuroscience Research (U01MH093765), part of the multi-institutional Human Connectome Project, and the Michael J. Fox Foundation for Parkinson’s Research (MJFF-021226). Computational resources were provided through the Massachusetts Life Sciences Center. B. Fischl is an advisor to DeepHealth, a company whose medical pursuits focus on medical imaging and measurement technologies. His interests were reviewed and are managed by Massachusetts General Hospital and Mass General Brigham in accordance with their conflict-of-interest policies.

